# Direct and indirect phenotypic effects on sociability indicate potential to evolve

**DOI:** 10.1101/2022.05.19.492047

**Authors:** David N. Fisher

## Abstract

The decision to leave or join a group is important as group size influences many aspects of organisms’ lives and their fitness. This tendency to socialise with others, sociability, should be influenced by genes carried by focal individuals (direct genetic effects) and by genes in partner individuals (indirect genetic effects), indicating the trait’s evolution could be slower or faster than expected. However, estimating these genetic parameters is difficult. Here, in a laboratory population of the cockroach *Blaptica dubia*, I estimate phenotypic parameters for sociability: repeatability (*R*) and repeatable influence (*RI*), that indicate whether direct and indirect genetic effects respectively are likely. I also estimate the interaction coefficient (*Ψ*), which quantifies how strongly a partner’s trait influences the phenotype of the focal individual and is key in models for the evolution of interacting phenotypes. Focal individuals were somewhat repeatable for sociability across a three-week period (*R* = 0.083), and partners also had marginally consistent effects on focal sociability (*RI* = 0.055). The interaction coefficient was non-zero, although in opposite sign for the sexes; males preferred to associate with larger individuals (*Ψ*_male_ = −0.129) while females preferred to associate with smaller individuals (*Ψ*_female_ = 0.068). Individual sociability was consistent between dyadic trials and in social networks of groups. These results provide phenotypic evidence that direct and indirect genetic effects have limited influence on sociability, with perhaps most evolutionary potential stemming from heritable effects of the body mass of partners. Sex-specific interaction coefficients may produce sexual conflict and the evolution of sexual dimorphism in social behaviour.

## Introduction

Many animals form groups and aggregations to find food, avoid predators, and to be buffered from environmental stressors (Krause & Ruxton, 2002). Individual sociability is therefore an important trait that can influence access to resources, mating opportunities, predators, and disease (Gartland *et al*., 2022). This importance means it is often linked to fitness. Further, in aggregate individual sociability determines group size, which in its own right can have influences on individuals’ fitness components (Silk, 2007). These links with fitness imply sociability is frequently under selection, and therefore would be expected to evolve if heritable. Predicting how individual sociability, and therefore also group size, will evolve requires us to estimate the genetic variance underpinning the trait i.e. its heritability (Scott *et al*., 2018). Typically, when estimating the heritability of a trait we consider its direct additive genetic variance i.e., how much variance among individuals in their own genes relates to variance in their phenotypes (hereafter direct genetic effects, “DGEs”). However, alongside its own social tendencies an individual’s sociability will likely depend on the traits of others in the groups it may join. For example, a normally sociable individual may be less willing to join a group with particularly aggressive individuals. As the traits of others will be at least partly influenced by genes, the heritable variation in sociability is likely to stem not only from DGEs, but also indirect genetic effects (IGEs), where genes in an interacting individual influence the focal individual’s trait (Griffing, 1967; Moore *et al*., 1997a). The presence of IGEs (and their covariance with DGEs) can accelerate evolutionary change, retard it, remove it completely, or even reverse it (Moore *et al*., 1997a; Wolf *et al*., 1998), potentially leading to non-linear responses to selection (Trubenová *et al*., 2015), responses to selection in the opposite direction to that of direct selection (Bijma & Wade, 2008; Fisher & Pruitt, 2019) and even maladaptation (Fisher & McAdam, 2019; McGlothlin & Fisher, 2021). Indirect genetic effects are widely appreciated in animal and plant breeding for their ability to prevent the evolution of higher yields (Muir, 2005; Ellen *et al*., 2014; Costa e Silva *et al*., 2017), and are becoming increasingly well appreciated in evolutionary ecology (Baud *et al*., 2022). If we want to understand how evolution shapes variation in sociability, the diversity of group sizes in nature, and how these traits might evolve in the future, we need to estimate how important both DGEs and IGEs are for individual sociability.

Despite the clear need to measure DGEs and IGEs on sociability, estimates of DGEs are quite rare (Lea *et al*., 2010; Brent *et al*., 2013; Staes *et al*., 2016; Knoll *et al*., 2018; Scott *et al*., 2018), and estimates of IGEs are completely absent (although Lea *et al*. did estimate DGEs for the tendency to *receive* interactions in a social network of marmots, which should be very similar to IGEs for initiating interactions). This can be partially attributed to two factors: 1) Experimental design to quantify individual sociability and how it is influenced by both direct and indirect effects can be difficult (Gartland *et al*., 2022) and 2) Estimating DGEs and IGEs in any context requires large amounts of both phenotypic data and information on genetic relatedness (Moore *et al*., 1997a; Bijma, 2014; Kruuk & Wilson, 2018). While 1) can be solved with appropriate experimental design, solving 2) can be logistically challenging. One partial (and temporary) solution is to estimate parameters that represent DGEs and IGEs at the phenotypic level, which does not require data on genetic relatedness and may also require less data overall as phenotypic variances are typically larger than genetic variances. Although not ideal, these parameters can still give insight into the evolutionary potential of the trait of interest as the relative magnitude of phenotypic and genetic variances (and covariances) are normally aligned (Hadfield *et al*., 2007; Dochtermann, 2011; Dochtermann *et al*., 2015).

For DGEs, the phenotypic parameter that (in most cases) sets the upper limit for heritability is repeatability (*R*, but see: Dohm, 2002). Repeatability is defined as the portion of phenotypic variance attributable to among individual differences (*VI*; Nakagawa & Schielzeth, 2010). This parameter can be decomposed into additive genetic variance and permanent environmental variance (*V_I_* = *V_A_ + V_PE_*), where for behavioural traits on average 52% of *V_I_* stems from *V_A_* (Dochtermann *et al*., 2015). We can therefore think of *R* as a phenotypic proxy for DGEs (as well as providing useful information about the relative balance between among- and within-individual variation in the population). Regarding IGEs, an analogous phenotypic equivalent in dyadic interactions would be the variance attributed to the identity of the interaction partner (*V_S_*). We could then calculate “repeatable influence” (*RI*) as the portion of phenotypic variance in the focal individual’s trait attributable to the among partner differences. For interactions with multiple partners, *V_S_* is multiplied by the average number of interaction partners (Bijma, 2011), and so can lead to the variance attributable to social interactions being greater than the phenotypic variance (Bijma *et al*., 2007; see: Ellen *et al*., 2014 for empirical examples of the total hertiable variation of a trait being greater than the phenotypic variance, due to IGEs), which makes it less directly relatable to *R*. Estimating both *R* and *RI* should give us a good indication of the *relative* contribution of DGEs and IGEs to a trait, while also giving some indication of the likely *absolute* magnitude of these parameters.

Alternatively to estimating DGEs and IGEs, a parameter widely used to infer their importance of social interactions in evolution is the interaction coefficient (*Ψ*; Moore *et al*., 1997a; Bailey & Desjonquères, 2022). This term is the coefficient from a regression of the focal individual’s phenotype on an interacting individual’s trait. It therefore does not require data on genetic relatedness. The *Ψ* term is key in the “trait-based” approach for understanding the role of social interactions in evolution, as opposed to the “variance-based” approach, which relies on DGEs, IGEs, and their covariance (McGlothlin & Brodie, 2009). *Ψ* can alter the direction and steepness of evolutionary trajectories, lead to feedback between interacting traits, and result in non-linear change (Bailey & Desjonquères, 2022). Additionally, *Ψ* can be converted into a direct-indirect covariance if the genetic variances of the traits of interest are known (McGlothlin & Brodie, 2009). We can therefore think of *Ψ* as both an important evolutionary parameter in its own right and as a phenotypic indicator of the likely magnitude of key genetic covariances. Together, *R, RI*, and *Ψ* give us useful gauges of the likely importance of DGEs and IGEs for a trait’s evolution, and so estimating them for sociability will give us a reasonable indicator for how this trait, and therefore group size, may evolve in the absence of genetic information.

Here, I estimated *R, RI*, and *Ψ* for sociability in the gregarious cockroach *Blaptica dubia* (Blattodea: Blaberidae). This is a communally living species who form aggregations in refuges to access resources, avoid predators such as ants, and to buffer environmental perturbations (Grandcolas, 1998) – hence their sociability is an important trait for their survival and fitness. Further, cockroach species are often sources of zoonoses (Zarchi & Vatani, 2009; Patel *et al*., 2022), and so a better understanding of the evolution of their aggregation formation could inform how the disease risk to humans they pose changes in the future under selection pressures such as changing climates and different management strategies. I measured sociability in dyadic trials and validated this assay in a group context using social networks of up to 24 individuals. I assayed individuals in the dyadic trials repeatedly to allow me to estimate consistency in sociability (*R*) and to isolate the consistent effect of a partner individual on the focal (*RI*). I also tested how a trait of the interaction partner influences the focal individual’s sociability to quantify *Ψ*. I used body mass as the trait in interacting individuals as it is typically heritable; Clark and Moore (1995) estimated the full-sibling heritability (likely to be an overestimate) of body mass in the Madagascar hissing cockroach (*Gramphadorhina portentosa*), which like *B. dubia* is in the Blaberidae family, as 0.93, while Moore et al. (2004) estimated the heritability of pronotum width in the speckled cockroach (*Nauphoeta cinerea*, also a Blaberid) as 0.62. Therefore, a clear estimate of *Ψ* for body mass would indicate social interactions are likely to be important for the evolution of sociability. I predicted that 1) there would be a correlation between the measures of sociability in the dyadic and group context, 2) that sociability will be repeatable, 3) that sociability will be repeatably influenced by the identity of the partner individual, and 4) that individuals will prefer to interact with larger partners (as smaller values in my sociability assay indicate more sociable, this means *Ψ* < 0) as they represent better protection from predators and the elements.

## Methods

### Experimental animals

*Blaptica dubia* is a quite large (up to 45 mm in length) sexually dimorphic blaberid cockroach (Wu, 2013). They typically live in aggregations at high temperature and humidity in central and south America (Alamer & Hoffmann, 2014), consuming vegetative matter, and are ovoviviparous. They are described as “gregarious” (Grandcolas, 1998) or “communal” as individuals of the same generation cohabit (without shared parental care; Bewick *et al*., 2017). I purchased an initial colony of *B. dubia* online in March 2021. I maintained them at the University of XXX [removed to preserve anonymity] at 28°C, 50% humidity, with a 50:50 light:dark cycle. I provided them with cardboard egg trays for shelter, carrot for hydration, and Sainsbury’s Complete Nutrition Adult Small Dog Dry Dog Food (approx, nutritional composition = 1527 kJ energy, 24g protein, 12g fat per 100g) for nutrition. Mortality was very low at all life stages (0.31% of the adults in the colony died per half week) indicating the colony was healthy. I moved newly born nymphs every few days to a container of dimensions 610 x 402 x 315 mm of similar aged individuals (density ranged from a few hundred of the earliest instars to 10-80 of later instars) and maintained them in mixed groups until adulthood (seven instars which takes approx. 250 days at this temperature; Wu, 2013). Upon reaching adulthood I moved them to either single sex groups (again in containers of 610 x 402 x 315 mm) or in small groups of two males and four to eight females in a container of dimensions 340 x 200 x 125 mm for breeding to maintain the stock population. For this experiment I selected 48 unmated males and 48 unmated females from the single-sex adult groups. Individuals were all more than five days old, and so presumed to be sexually mature (Hintze-Podufal & Vetter, 1996). I transferred each individual to a clear plastic box (79 x 47 x 22 mm) labelled with its unique ID to allow individual recognition. I gave individuals a small piece of carrot for hydration which was replaced weekly.

### Data collection

I tested individuals in two blocks of 48, treating all individuals in each block once as a focal individual and once as a partner for a member of the same sex over two days. This means that in the first two days 24 males and 24 females were each assayed for sociability once and each acted as a partner individual once. On days three and four I repeated this with a second block of 24 males and 24 females. In this way individuals only ever acted as focal or partner individuals with members of the same sex in the same block (either first or second) and were each assayed for sociability and acted as a partner once per week. I repeated this for three weeks, so each individual was assayed up to three times as a focal individual as acted as a partner up to three times. Some individuals received fewer than three trials if they died (n = 5 males and 0 females), in which case I replaced them with a member of the same sex from the stock population (who did *not* inherit the same ID and was therefore another unique individual). Individuals might also record fewer than three measures for sociability if the mesh was breached by either the partner or the focal before the trial began (11 females and eight males recorded one measure, 48 females and 42 males recorded two measures, 30 females and 54 males recorded three measures).

I assayed sociability in medium sized plastic boxes (200 x 100 x 70 mm) where I glued a fine polypropylene mesh (mesh size 0.6 x 0.6 mm, Micromesh, Haxnicks) across the interior 50 mm from one end. This creates an arena with a small compartment (50 x 100 x 70 mm) and a large compartment (150 x 100 x 70 mm) separated by the mesh (Fig. 1A). Separating by mesh was necessary to prevent a partner individual imposing close proximity on the focal individual by constantly following or attempting to dominate it (Clark *et al*., 1995), and therefore my assay captures the focal individual’s willingness to socialise, rather than the partner’s (Gartland *et al*., 2022). For the first block I randomly placed 12 females and 12 males alone, each into their own plastic box, in the large compartment. These were the focal individuals. I then randomly placed an individual of the same sex into the small compartment; these were the partner individuals. I used individuals of the same sex to ensure I was measuring sociability rather than willingness to mate. I then placed these 24 arenas into four large plastic boxes (six in each) which I placed underneath a video camera (ABUS IP video surveillance 8MPx mini tube camera), so that each video camera recorded six arenas simultaneously. I maintained the room the video recordings occurred in at 20-22°C using portable heaters, while I used a thermometer to record the temperature at the start and end of each trial. I was not able to control or monitor humidity during the trials. Once all arenas were in position and cameras focused, I started the recording and left the room. The lights automatically switched off after 40 minutes, and so the trial began 40 minutes after I left the room, in darkness, which is when *B. dubia* is active (Bouchebti *et al*., 2022). I returned two hours after leaving to end the trial, meaning the trials lasted 80 minutes. In darkness the cameras automatically switch to infra-red filming using infra-red LEDs.

**Figure 1.**
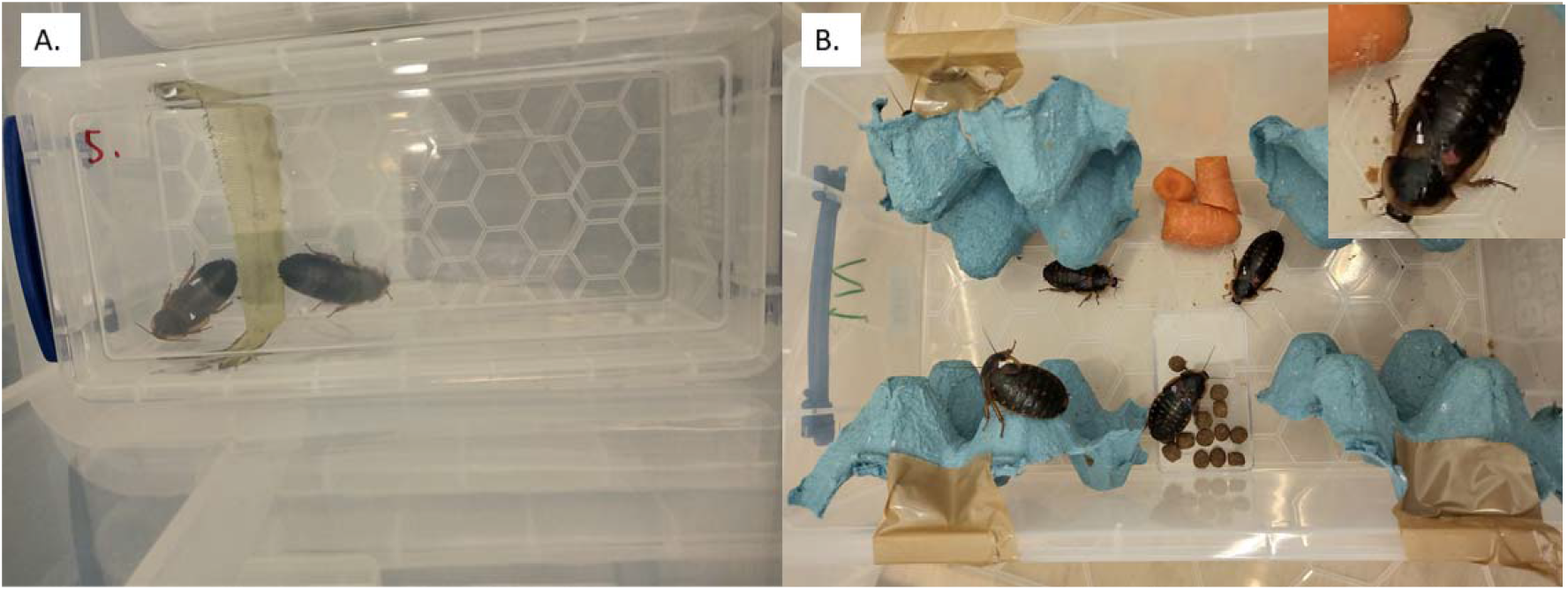
Pictures of experimental set-up (both XX [removed to preserve anonymity]). A. Assay for sociability. The position of the focal individual (on the right in the larger compartment, no white dot) in relation to the mesh is recorded every ten minutes to assess willingness to socialise. This individual would record a score of two. B. Social network trials. Marked individuals (here showing green-blue, red-white, white-blue, and green-white, starting at the top left and moving clockwise) can chose among four equal shelters (the cardboard egg trays taped to the sides of the box). Co-occurring at a shelter with the same individual regularly indicates a social association. A close up of red-white is shown in the upper right corner.

For each trial, every ten minutes I recorded the proximity of the focal individual (in the larger compartment) to the mesh that separated it from the partner individual (Fig. 1A), giving a maximum of eight measures per trial. The distance of an individual to a conspecific in this manner is often used to measure sociability (reviewed in: Gartland *et al*., 2022) and by using the distance to the mesh rather than the partner I have a measure solely under the influence of the focal individual; the partner cannot directly influence the distance by moving closer or further away. If the focal was sat directly on the mesh (perpendicular to the floor) I recorded a distance of zero, otherwise I used the hexagons on the bottom of the box to record how far the focal individual’s head was from the mesh (Fig. 1A). Smaller values mean a focal individual closer to the partner individual which indicates higher sociability. Individuals were in some cases able to bypass the mesh (this occurred 95 times before the lights went out and 33 times after they did out of 288 trials, the 33 breaches after lights out are still included in the analyses with only the measurements before the breach used, see Data analysis). I used the video recordings to determine when this happened and stopped recording data from the video as soon as either individual bypassed the mesh. If either individual bypassed the mesh in the 40 minutes the lights were on before the trial started then I recorded no data from that trial. To avoid mixing individuals up at the end of the trial I dotted either the partner or the focal with white paint (Edding Extra-fine paint markers). At the end of the trial, I returned all individuals to their unique boxes. I then weighed all partner individuals to the nearest 0.01 g (Fisherbrand Analytical Balances, readability 0.0001 g). In the first two days I also weighed the focal individual, but since the correlation between an individual’s mass as a focal and its mass as a partner either the following or preceding day was 0.994 (Pearson correlation, t_38_ = 56.894, p < 0.001) I stopped doing this to save time. Instead, I entered the mass of the focal individual as its mass as a partner individual recorded the same week (always within one day). As described above, each individual in the two blocks was assayed once as a focal and acted once as a partner for another individual of the same sex in that block per week, and this was repeated for three weeks.

After the third trial I aggregated individuals into four groups of 21-24; all the individuals of the same sex from the same block were together, with groups having fewer than 24 individuals if any died (I did not replace individuals that died with stock individuals as I was only interested in the social network position of those with a known sociability from the dyadic trials). I gave each individual a unique combination of two colours (red, green, blue, white, gold) on their wing cases using paint pens (Edding Extra-fine paint markers; Fig. 1B) which allowed me to track them individually (combinations were repeated between groups i.e., red-blue featured in each of the four groups). I then placed each group into new plastic boxes (340 x 200 x 125 mm) along with four shelters made from cardboard egg tray (approx. 100 x 120 mm), each placed vertically at each corner on a long side (Fig. 1B). Shelters were taped to the walls of the box, creating clear space between both the shelter opposite it (on the opposite long side) and next to it (on the same long side). I placed 2 g dog food and 10 g carrot in the centre of each box. Each shelter was large enough to accommodate many but not all of the individuals, and the number of shelters was considerably less than the number of individuals. Therefore, the formation of aggregations in shelters was enforced, but individuals could move between shelters and therefore could exert some influence on who they co-habited with. Regularly after placing the individuals into these groups (after 3, 10, 14, 18, and 21 days) I recorded which individuals were using the same shelter. Individuals who could not be identified were recorded as such but they were not used to build the networks. Collecting data in this way gives a group-by-individual matrix analogous to those generated by observing flocks of birds or herds of ungulates in the wild, and further is similar to methods than have been used to generate social networks in forked fungus beetles (*Bolitotherus cornutus*; Formica *et al*., 2012, 2016, 2020) and maritime earwigs (*Anisolabis maritima*; Vipperman, 2021). While a single incidence of sharing a shelter could be due to chance, by aggregating these observations I can infer consistent social associations. When recording these data, I also updated any paint markings that were starting to wear, maintaining individually-recognisable marks for the duration of the experiment, and replaced carrot and dog food as necessary.

### Data analysis

All analyses were conducted in R (version 4.1.3; R Development Core Team, 2016). To analyse sociability, I summed each individuals’ distances from the mesh across the 1-8 records per trial and entered that as a response variable in a generalised linear mixed effects model using *glmmTMB* (Brooks *et al*., 2017). To account for the different number of measures contributing to this sum (if individuals “breached” the barrier during the trial) I included an offset of the log of the number of records the individual recorded from the trial and used a Poisson error distribution and a log link function. While the response variable entered into the model is still the sum of the distances (and hence an integer), this approach effectively models the mean distance the individual is from the mesh (sum / n. trials) but is preferable from directly using this variable as it can be used with a Poisson error distribution, which requires integers and so is incompatible with the mean distance (the residuals are also greatly improved, see Fig. S1). I included fixed effects of the temperature in the room (scaled to a mean of zero and a standard deviation of one), the sex of the individual (and therefore also its partner), the body mass of the focal individual and the body mass of the partner, both scaled to a mean of zero and a standard deviation of one, and the interaction between both focal and partner (scaled) body mass and sex. The effect of the partner mass is key for testing prediction 4 as its coefficient is our (unstandardised) estimate of *Ψ*, while the interaction with sex tests whether this differs between the sexes. I included random effects of individual ID, partner ID, and date, to estimate the variance among focal individuals, partner individuals, and dates respectively. To estimate *R* for sociability (testing prediction 2) I extracted the model intercept, the among-focal individual variance, and the sum of all variance components, and entered them into the ‘QGicc’ function in the package *QGglmm* (de Villemereuil *et al*., 2016), using the ‘model = “Poisson.log”’ setting. This calculates *R* for sociability on the original scale as opposed to the latent scale (Nakagawa & Schielzeth, 2010; de Villemereuil *et al*., 2016); the former is necessary to compare to estimates of *R* from traits analysed assuming a Gaussian distribution. I repeated this with the among-partner individual variance instead of the among-focal individual variance to obtain the estimate of *RI* (testing prediction 3). Alongside the magnitudes of *R*, I demonstrated the importance of accounting for differences among individuals in sociability by comparing the AIC of the model described above to a model identical except that the random effect of focal individual was removed. I did the same for *RI* i.e., comparing models with and without the partner ID term (the models were otherwise identical to the one described above). To determine the clarity of fixed effects I used the ‘Anova’ function in the package *car* (Fox & Weisberg, 2019) with a type three sum of squares to generate p values (see: Dushoff *et al*., 2019 for a discussion on the use of “clarity” over “significance”).

After finding a clear interaction between body mass and sex (see Results), I wished to obtain sexspecific estimates of *Ψ* that were standardised to facilitate comparisons across studies (note this was a decision made after viewing the initial results and so should be interpreted more cautiously). To do this I refitted the above model to the sexes separately (minus the fixed effect of sex and its interactions with focal and partner body mass), this time with the mean distance the focal individual was from the mesh, scaled to a mean of zero and a standard deviation of one, as the response variable, with no offset and assuming a Gaussian error distribution with a normal link function. Using this transformed variable generates estimates of *Ψ* that are comparable across traits and studies (Bailey & Desjonquères, 2022).

To generate a social network per group I used each groups’ group-by-individual matrix, which contains records of which individual was in which shelter at each time point. From this I created four networks where individuals were linked with their relative association strengths, which is undirected i.e., individual A’s interaction with individual B is equal to individual B’s interaction with individual A. I calculated relative associations strengths as the simple ratio index (Cairns & Schwager, 1987), using the package *asnipe* (Farine, 2013). This index is the count of all times individuals shared a shelter, divided by the number of occasions both individuals were recorded (this could be less than five if an individual died during this phase of the study), and so indicates the relative strength of the association between any two individuals. A score of one indicates two individuals who were always seen sharing a shelter, and zero two individuals who never shared the same shelter. I summed each individual’s association scores to gives that individual’s ‘strength’, a measure of network centrality that captures an individual’s overall engagement in social interactions and in this case is analogous to the average group size an individual was found in.

To test whether sociability as determined by the dyadic assays correlated with sociability in the social network (prediction 1), I followed the suggestions of Hadfield et al. (2010) in the guide from Houslay and Wilson (2017) to estimate the among individual correlation between the two traits (see also: Dingemanse *et al*., 2012; Dingemanse & Dochtermann, 2013). This approach excludes the residual variance from the correlation, specifically addressing our question of interest (are individuals that are more sociable in the dyadic assay more sociable in the social network). To do this I fitted a bivariate mixed-effects model in MCMCglmm (Hadfield, 2010). The response variables were each of an individual’s sum of locations from the dyadic trials, and its strength as quantified in the social network. As there is only one value of the latter it is repeated each time an individual records a sociability score i.e., 1-3 times. I included the fixed effect of the log of the number of observations and modified the prior to constrain the relationship between this fixed effect and the sum of locations or network strength to 1 or 0 respectively (by setting the coefficients to 1 and 0 respectively and setting both variances as 1 x 10^-9^). This approach is equivalent to fitting number of observations as an offset for the sum of locations, and as having no relationship at all with network strength. The random effect was a 2 x 2 covariance matrix estimating the among individual variance in each trait and the among-individual covariance between them (our parameter of interest). I allowed the residual variance for sociability in the dyadic assay to be non-zero (as there are multiple measures on individuals) while I fixed it at 0.0001 for network strength (as there is only a single measure it does not vary within individuals), and I fixed the residual covariance at zero. I set a Poisson error distribution for sociability in the dyadic assay and a Gaussian error distribution for network strength. I used 550,000 iterations, with the first 50,000 discarded and 1 in 100 of each subsequent iteration retained. I confirmed the model had converged by running three chains and calculating the Gelman and Rubin convergence diagnostic (Gelman & Rubin, 1992; point estimates were all 1.01 or lower), as well as assessing the trace plots. I calculated the among individual correlation between the two measures of sociability as the among-individual covariance between the two traits divided by the square root of the product of their two among-individual variances and extracted the mode and 95% credible intervals from the resulting posterior distribution.

## Results

I collected 193 observations of sociability across 92 individuals in dyadic trials, and quantified the social network position of each of these individuals (Fig. 2). Individuals that were more sociable in the pairwise trials had higher strength in the social network trials (Among-individual correlation between sociability and network strength, posterior distribution mode = −0.758, 95% credible intervals = −0.948 to −0.265), confirming prediction 1 and validating the use of the dyadic trials to assay sociability. Full model results are shown in Table S1.

**Figure 2.**
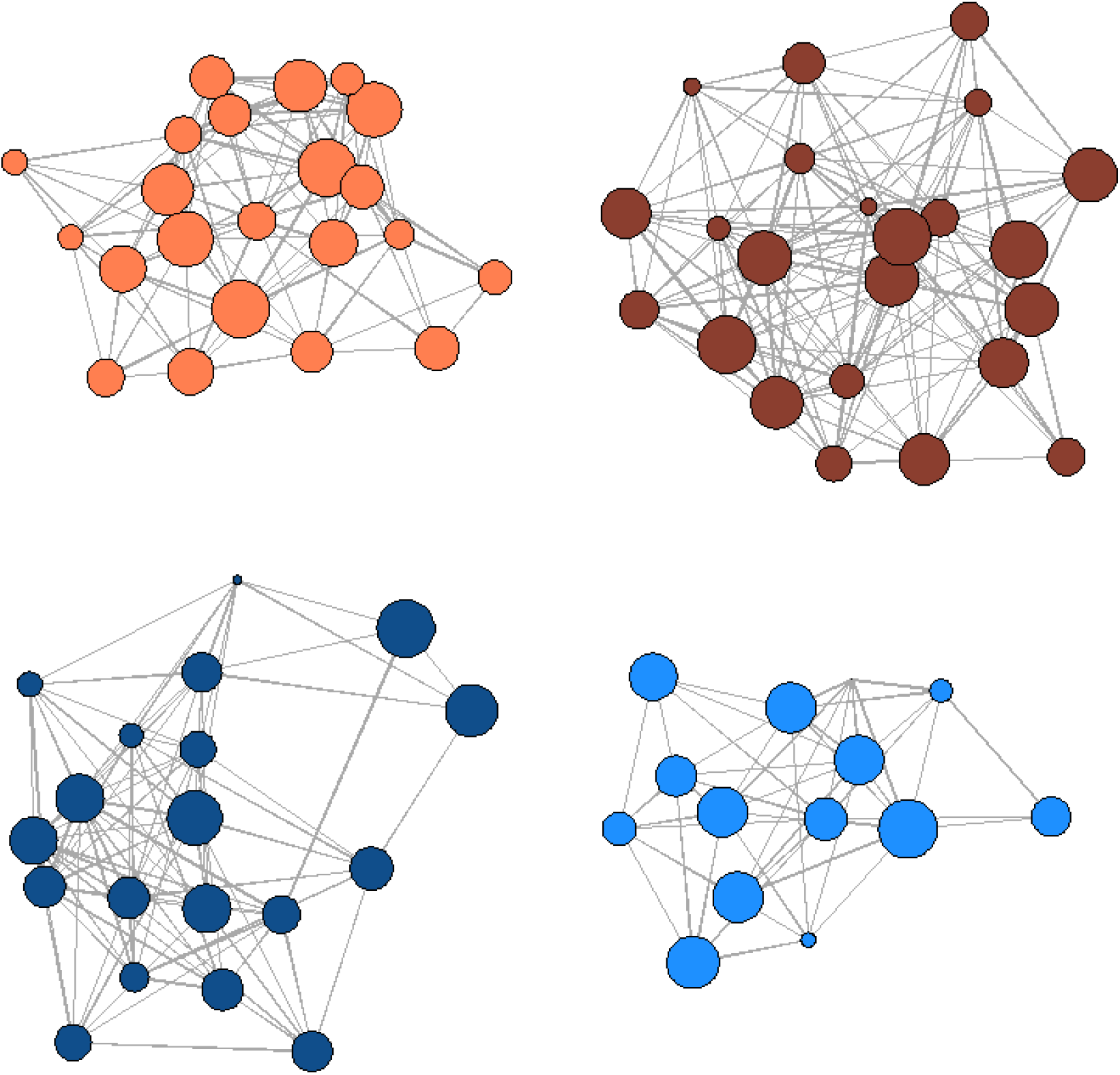
Social networks of *Blaptica dubia* individuals for each group (females on top row, males on bottom row, node size indicates sociability in pairwise trial [larger = more sociable], thickness of lines indicates strength of association). I have removed edge weights below 0.12 to reduce visual clutter.

Individuals are weakly repeatable in how sociable they are, with an *R* of 0.083 (prediction 2), while the ΔAIC of model with vs without the random effect of focal individual ID was 767. Individuals exerted a small amount of repeatable influence on the sociability of their partners, with an *RI* of 0.055 (ΔAIC = 774; prediction 3). The body mass of the partner individual influenced how sociable the focal individual was (prediction 4), with males being more sociable with larger individuals and females being more sociable with smaller individuals (Fig. 3; main effect β = 1.350, se = 0.463, χ2 = 8.493, p = 0.004, interaction β = −2.833, se = 0.686, χ2 = 17.038, p < 0.001). Full model results are presented in Table 1. Given this clear interaction, I decided post-hoc to fit separate models to each sex to generate sex-specific and standardised estimates of *Ψ*, giving *Ψ*_female_ = 0.068 (se = 0.103) and *Ψ*_male_ = −0.129 (se = 0.107; recall that lower scores in the pairwise trial indicate higher sociability as they represent shorter distances from the partner individual). There are therefore phenotypic indicators that there are IGEs on individual sociability, but the direction of the effect is opposite in sign for the sexes and both are quite near zero.

**Figure 3.**
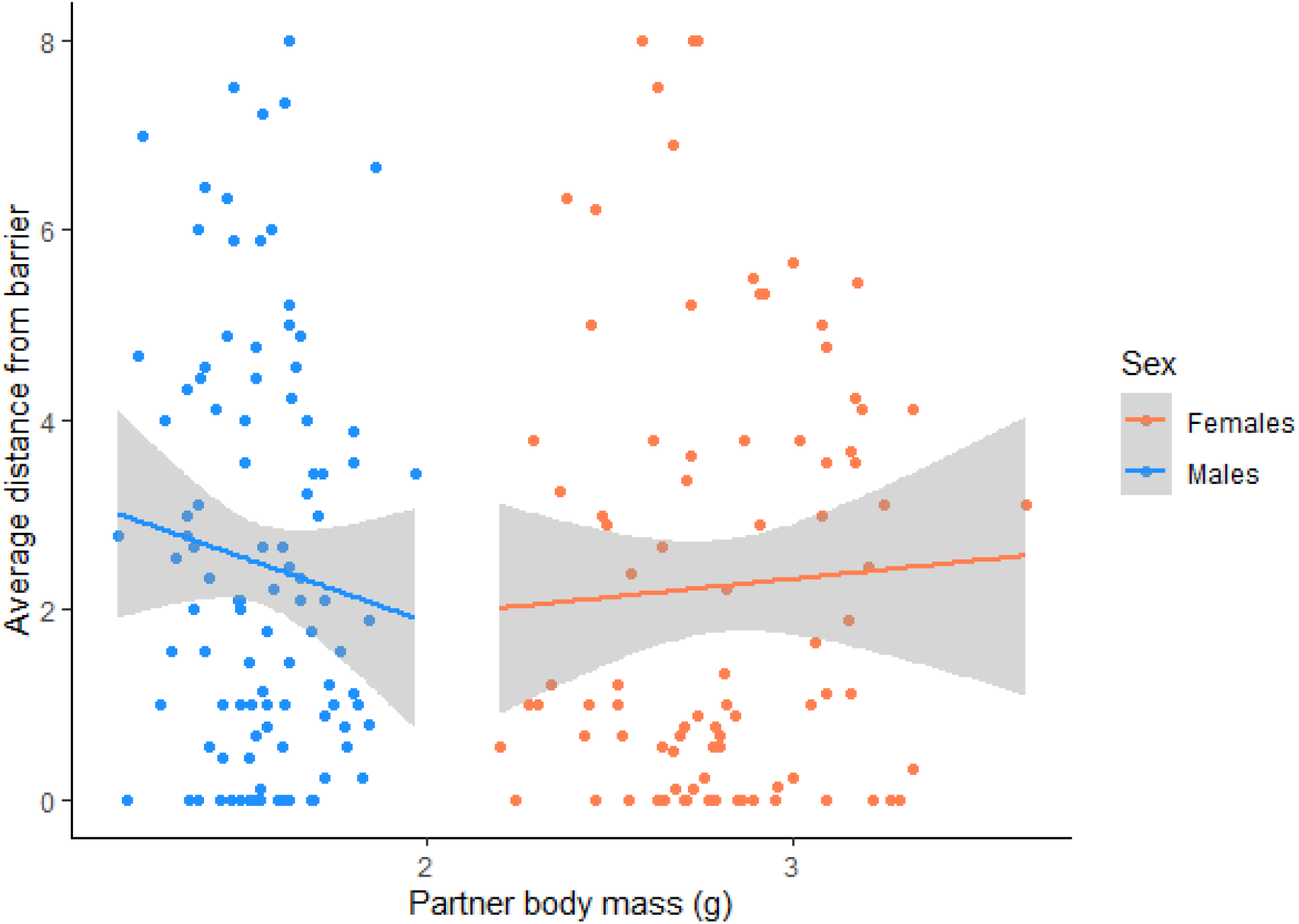
The body mass of the partner individual influences the sociability of the focal individual, with males (blue) preferring to be near heavier individuals, while females (orange) preferring to be nearer lighter individuals. Points are individual scores while lines indicate the mean effect estimated by the plotting function. Grey areas indicate the 95% confidence intervals around the mean.

**Table 1.** Full model output for analysis of sociability in dyadic trials. Females were set as the default sex and so the effect of sex is the contrast between males and females.

| Variable | Estimate | Standard Error | Chi-squared | P value |
| --- | --- | --- | --- | --- |
| Intercept | -0.476 | 0.584 | 0.665 | 0.415 |
| Sex (Male contrast) | -1.787 | 0.986 | 3.283 | 0.070 |
| Body mass of focal | -0.940 | 0.450 | 4.359 | 0.037 |
| Body mass of partner | 1.350 | 0.463 | 8.493 | 0.004 |
| Temperature | 0.168 | 0.178 | 0.897 | 0.343 |
| Sex : Body mass of focal | -0.345 | 0.586 | 0.347 | 0.556 |
| Sex : Body mass of partner | -2.833 | 0.686 | 17.038 | 0.000 |
| Random effect |  |  | Variance |  |
| Individual |  |  | 2.503 |  |
| Partner |  |  | 2.130 |  |
| Date |  |  | 0.281 |  |

## Discussion

I found that the measure of sociability in the dyadic trials was correlated among-individuals with a measure of centrality from a social network, validating the use of the dyadic trials for assaying sociability. Sociability in the dyadic trials was repeatable, and partners had a repeatable influence on the sociability of others, but both values were small. This implies that the direct and indirect genetic variance in the trait is likely to be low, and hence selection on this trait will be converted into limited evolutionary change. I also found the body mass of the partner influenced the sociability of the focal individual, but in sex-specific ways; males were more sociable with large individuals, while females were sociable with small individuals. As body mass is typically heritable this suggests that there is some evolutionary potential in sociability through the effect of body size; if body mass evolves, so will sociability, but in opposite directions for males and females.

Finding sociability had non-zero repeatability is to be somewhat expected as behaviours typically are repeatable (Bell *et al*., 2009; Holtmann *et al*., 2017). Further, a low repeatability might be expected given social behaviours often depend on the phenotype of interaction partners which can be expected to vary substantially in short periods of time (Holtmann *et al*., 2017). Further, I found relatively little variance attributed to the identity of the partner individual. To explore this further, I re-ran the model with the mass of the partner (and its interaction with sex) removed. This reduced model gave a slightly higher estimate of *V_S_*(2.19 vs. 2.13). Therefore, while body mass exerts some consistent effect on other individuals, other traits also contribute. In any case, most of the variation in sociability was not explained by the variance components or fixed effects, indicating sociability is highly labile.

Given that, on average, 52% of the among-individual variance of behaviours stems from direct additive genetic variance (Dochtermann *et al*., 2015), we can expect sociability to have a low heritability even when the indirect genetic variance is included. A low heritability means selection would be translated into a small amount of evolutionary change, and so sociability and therefore mean group size may show limited response to direct selection. In contrast, Dochtermann *et al*. (2019) used a meta-analysis across taxa to infer that social behaviours had heritabilities typical of behavioural traits; similar to the overall mean of 0.235. This estimate however only considered direct heritability and not indirect genetic effects, the total heritability may well be even higher (or lower, depending on whether the direct-indirect genetic covariance is positive or negative; Bijma, 2011). Estimating the direct and indirect heritability of sociability and obtaining ecologically relevant estimates of selection (or relationships between sociability and fitness components such as survival or longevity; Blumstein *et al*., 2018; Brodin *et al*., 2019; Montero *et al*., 2020) are logical next steps to better understand the microevolution of this trait (see also the artificial selection experiment of Scott *et al*., 2022 who successful increased sociability in *Drosophila melanogaster* over 25 generations).

Alongside the repeatability of sociability, I found that how sociable an individual was depended on its own mass and the mass of the mass of the partner individual. While heavier individuals of both sexes were more sociable, females preferred to be near smaller females, while males preferred to be near larger males i.e., *Ψ* was sex specific. While I predicted an effect of partner mass, I had not predicted a sex-specific effect. Females are larger than males and require protein for egg production which males do not (Maklakov *et al*., 2008; Jensen *et al*., 2015), and so females may be in more intense competition for resources with each other than males are. This competition could lead to them preferring to associate with smaller individuals who are presumably less competitive. In contrast, males may prefer larger individuals as they offer better protection from predators, more protection from desiccation, and possibly if larger males attract females, the so called “hotshot” effect (Beehler & Foster, 1988). Alternatively, both sexes may be seeking mating partners (I used unmated individuals), which are always smaller or larger than themselves for females and males respectively (see how the masses of the sexes do not overlap in Fig. 3). This would lead to females preferring to be near smaller females who are perhaps harder to distinguish from males, and vice versa for males (suggested by Han *et al*., 2016, although they found no effect of partner body size on same-sex behaviour in water striders *Gerris lacustris*). While we would expect chemical communication to be important for mate choice cockroaches (Schal *et al*., 1984; as well as for other social interactions: Moore, 1997; Moore *et al*., 1997b), it is possible individuals use both chemical cues and morphological traits when searching for a partner. Testing these ideas, and the fitness consequences for both males and females for associating with large and small individuals (“social selection”; Wolf *et al*., 1999; e.g.: Santostefano *et al*., 2019; Fisher *et al*., 2021), represent key next steps. Furthermore, it is worth highlighting that body mass represents only one trait that influences sociability; there may well be other morphological, behavioural, or chemical traits of partners that affect focal individual behaviour. While these are accounted for in the estimate of *V_S_*, identifying the causal traits is a useful step for understanding the mechanisms underpinning social interactions.

Indirect genetic effects can fundamentally alter the direction and magnitude of evolutionary change, and so finding opposing estimates of *Ψ* for the sexes implies that sociability in the sexes could follow quite different evolutionary trajectories. Whether they will do so or not depends on the genetic variance underpinning mass (which is likely to be non-zero, see Introduction) and sociability and the inter-sex genetic correlation for sociability (McGlothlin & Brodie, 2009). Further, given preferences differ, selection on partner choice may well differ between the sexes (Parker, 1979; “sexually discordant selection”; Westneat & Sih, 2009). Such selection can be expected when sexual dimorphism cannot evolve, possibly due to inter-sex correlations typically being large and positive (Poissant *et al*., 2010). In the case of *B. dubia*, any quantitative predictions at this stage would be premature given the number of assumptions I would be required to make, but it is interesting that same-sex social interactions potentially facilitate sexual conflict thanks to estimates of *Ψ* which are opposite for the sexes. Results here show that if population mean body mass increases, male sociability will increase, while female sociability will decrease. In the short term at least (before female preferences change) reduced sociability could reduce female mating rate, influencing the costs and benefits of polyandry such as male harassment, sexual infection transmission, direct and indirect benefits from mating with multiple males. In general, however, the estimates of *Ψ* (which typically range from −1 to 1; Bailey & Desjonquères, 2022) in this study are quite near zero, indicating only modest deviations from a situation where individuals do not impact each other’s traits through social interactions (this is quite common for estimates of Ψ between different traits; Bailey & Desjonquères, 2022).

My results add to increasing evidence that *Ψ* varies among- (Kent *et al*., 2008; Bleakley & Brodie III, 2009; Bailey & Zuk, 2012; Edenbrow *et al*., 2017; Marie-Orleach *et al*., 2017; Culumber *et al*., 2018; Kraft *et al*., 2018) and within-populations (Edenbrow *et al*., 2017; Signor *et al*., 2017; Han *et al*., 2018). *Ψ* itself can evolve (Chenoweth *et al*., 2010; Bailey & Zuk, 2012; Rebar *et al*., 2020), and the evolution of *Ψ* can both increase or decrease the speed of evolutionary change (Kazancio⍰lu *et al*., 2012). It is therefore clearly important to study this parameter in an evolutionary context to better understand the evolution of interacting phenotypes (Bailey & Desjonquères, 2022). This is especially true when the majority of the evolution potential of a trait may stem not from genetic variance in the trait (which is what is typically assumed), but from associations with other heritable traits (such as body mass in this study). Additionally, it is especially important to estimate *Ψ* in systems where is expected to be large. Bailey & Desjonquères (2022) highlight that large and positive values of *Ψ* occur when individuals interact through the same trait, suggesting a positive feedback loop in trait expression. This would mean small initial evolved changes in a trait would lead to increased expression of the same trait in the partner, and so the trait mean would increase rapidly (Moore *et al*., 1997a). High and positive values of *Ψ* are observed in shoaling fish (Poecilia reticulata; Bleakley & Brodie III, 2009; Marie-Orleach *et al*., 2017; Gambusia holbrooki; Kraft *et al*., 2018), fruit flies (*Drosophila melanogaster* and *D. simulans;* Bailey & Hoskins, 2014; Signor *et al*., 2017) and burying beetles (*Nicrophorus vespilloides;* Rebar *et al*., 2020). More estimates of *Ψ* need to be accumulated before more general patterns in when *Ψ* strongly influences evolutionary potential can be identified (Bailey & Desjonquères, 2022).

The sociability of an individual estimated through repeated pairwise trials over three weeks was related to the individual’s centrality, in terms of its overall number and strength of associations, in a social network formed over 21 days. This result indicates that my assay for sociability accurately captures a facet of individual social behaviour, and this social behaviour is trait of an individual that is consistent across time and across contexts, and hence could be associated with lifetime reproductive success (Kluen & Brommer, 2013; although this association cannot be very strong as the trait’s repeatability is low). What maintains consistent among-individual differences in sociability and social network position are open questions (Wilson *et al*., 2012; Wilson & Krause, 2014; Gartland *et al*., 2022). If the behaviour is indeed heritable, then different levels of sociability should give similar fitness on average, as otherwise selection would remove the variation in sociability from the population. Different levels of sociability may therefore represent different strategies that bring both benefits (e.g., a higher sociability decreases water loss) and costs (e.g., a higher sociability decreases access to resources). Furthermore, switching between strategies must impose costs in some way so that individuals cannot be completely plastic (Dall *et al*., 2004; Snell-Rood, 2013). A second, not mutually exclusive, mechanism that could maintain variation in social strategies is if selection on the strategy is negative frequency dependent (Bergmüller & Taborsky, 2010). Better understanding of patterns of selection for this and similar traits is therefore key for predicting evolutionary responses in natural populations, and, ultimately, population dynamics and viability.

In summary, sociability in *B. dubia* shows a small degree of repeatability, some consistent influence from the identity of a partner, and is correlated among-individuals between trials in pairs and trials in groups. Males preferred to associate with larger individuals while females preferred to associate with smaller individuals. The latter result suggests that the evolution of sociability, and therefore the evolution of group size, may fundamentally depend on evolutionary change in body mass, and could drive sexual dimorphism in social behaviour. These sex-specific estimates of *Ψ* will be important for informing our models predicting microevolutionary change and for understanding sexual conflict. Future work will need to assess the fitness consequences of social behaviour and identifying the factors that predict patterns of social interactions in various more ecologically relevant settings.

## Acknowledgements

I would like to thank Keith Lockhart for his invaluable work maintaining the stock population. Maria Moiron and Francesca Santostefano provided numerous useful comments and suggestions on an earlier draft. Two anonymous reviewers made constructive comments. Funding was provided by the University of Aberdeen “Internal Funding to Pump-Prime Interdisciplinary Research and Impact Activities” fund. I have no conflicts of interest.

## Data accessibility

R code to conduct the analyses in this manuscript can be accessed at https://github.com/DFofFreedom/Direct-and-indirect-phenotypic-effects-on-sociability-. The associated data will be added upon acceptance.

