## Supplementary materials for "Direct and indirect phenotypic effects on sociability indicate potential to evolve"

Supplementary Tables

**Table S1**. Full results of the multivariate model between pairwise sociability and network strength. Given are the variable names, whether they are fixed or random effects, or were a correlation calculated from the model results, the posterior distribution mean and mode, the lower and upper 95% credible intervals (CrI) for the posterior distribution, the effective sample size for the variable, and the pMCMC value for the fixed effects. Variable names with an asterisk were fixed and so not estimated. Model fitted in MCMCglmm (Hadfield 2010), table produced using the package stargazer (Hlavac 2022) and code from https://gkhajduk.github.io/2017-10-25-cleanMCMCglmm/ .

| Variable | Effect | Posterior Mode | Posterior Mean | Lower 95% CrI | Upper 95% CrI | Effective sample size | pMCMC |
| --- | --- | --- | --- | --- | --- | --- | --- |
| Pairwise sociability Intercept | fixed | 0.154 | 0.156 | -0.072 | 0.380 | 5000.000 | 0.185 |
| Network strength Intercept | fixed | 4.001 | 3.980 | 3.704 | 4.231 | 400.186 | 0.000 |
| Pairwise sociability Offset* | fixed | 1.000 | 1.000 | 1.000 | 1.000 | 5355.755 | 0.000 |
| Network strength Offset* | fixed | -0.000 | -0.000 | -0.000 | 0.000 | 4634.383 | 0.994 |
| Pairwise sociability Individual ID | random | 0.117 | 0.222 | 0.000 | 0.498 | 2895.317 | - |
| Network strength Individual ID | random | 1.377 | 1.417 | 1.007 | 1.859 | 375.178 | - |
| Pairwise sociability – Network strength Among-individual covariance | random | -0.326 | -0.337 | -0.653 | -0.043 | 414.689 | - |
| Pairwise sociability – Network strength Among-individual correlation | correlation | -0.758 | -0.635 | -0.948 | -0.265 | - | - |
| Pairwise sociability Residual | residual | 1.801 | 1.824 | 1.320 | 2.381 | 5000.000 | - |
| Network strength Residual* | residual | 0.000 | 0.000 | 0.000 | 0.000 | 0.000 | - |

Supplementary Figures

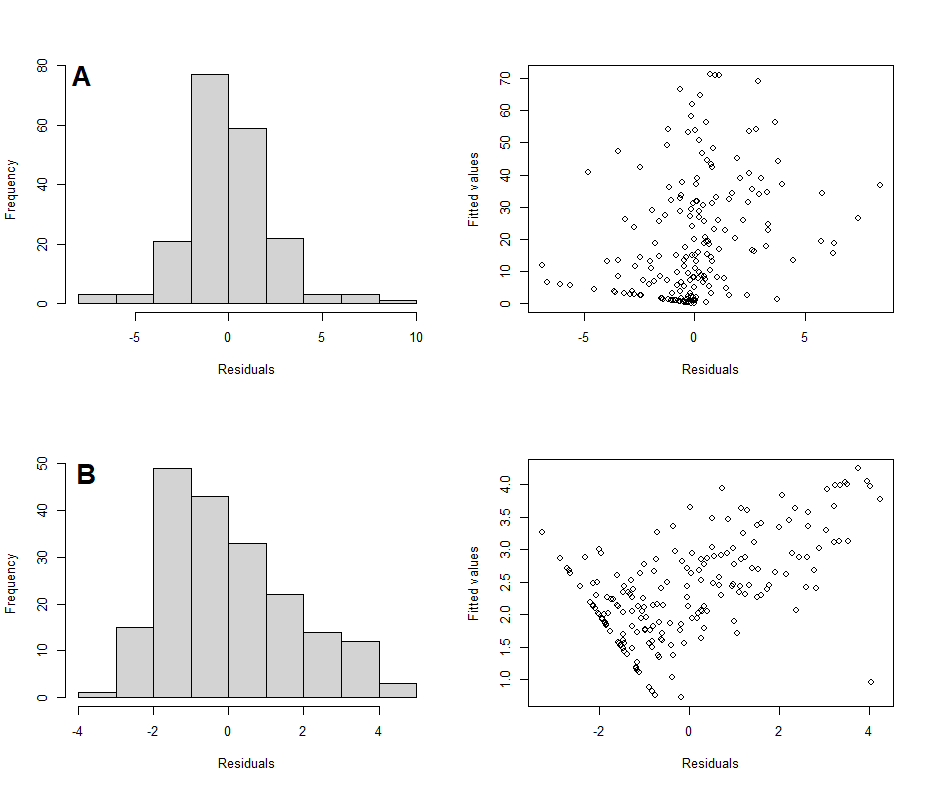

**Figure S1**. A comparison of the distribution of the residuals (left) and the relationship between residuals and fitted values (right) for the model fitted using the sum of locations and an offset (A) and a model using the mean location with no offset (B). Plots for A show more normally distributed residuals and no linear relationship between residuals and fitted values compared to plots for B.
